# Classification of Visual Imagery and Imagined Speech EEG based Brain Computer Interfaces using 1D Convolutional Neural Network

**DOI:** 10.1101/2024.12.11.627386

**Authors:** Diane Le

**Affiliations:** Department of Computing, Goldsmiths, University of London

## Abstract

Non-invasive brain-computer interfaces (BCI) utilising electroencephalogram (EEG) signals are a current popular, affordable and accessible method for establishing communication paths between the mind and external devices. However, the challenges faced are inter-subject variability, BCI illiteracy and poor machine learning decoding performance. Two emerging intuitive mental paradigms, Visual Imagery (VI) and Imagined Speech (IS) show promise to optimise the development of non-invasive BCIs, which involves the extraction of corresponding neural patterns during the imagined tasks. This study took a comprehensive user-centric approach to build on the current foundation of knowledge on VI and IS EEG-BCIs utilising an adapted 1D-CNN to optimise the classification decoding performance. Twenty healthy participants were assessed for their ability to visualise imagery in their minds and performed the VI and IS mental paradigms in two class conditions “push” and “relax”. It was shown that alpha and beta suppression was observed during the “push” condition of VI compared to the “relax” condition, and those that scored higher in the VVIQ had better VI classification accuracy than those who did not. The adapted 1D-CNN model performed well for classification between the two classes “push” and “relax” at 89.3% and 77.87% performance accuracy for VI and IS, respectively. These findings contribute to the current body of work on VI BCI, that it is a dynamic and plausible option compared to standard BCI paradigms, and VI BCI illiteracy could potentially be controlled by VVIQ. It also demonstrated the potential of the 1D-CNN model in classification of VI and IS EEG-BCIs.

## 1. Introduction

In the rapidly evolving landscape of neurotechnology, Brain-Computer Interfaces (BCIs) have emerged as a compelling solution, drawing attention for their potential to bridge the divide between the human mind and machines, especially in the use for patients with Locked-In-Syndrome (LIS) to communicate with the world. Fundamentally, BCIs enable the translation of neural signals into actionable commands that can be interpreted by computers or machines, thus unlocking the potential for direct brain-mediated control and interaction. However, the pursuit of refining and expanding non-invasive BCI applications has spurred the exploration of novel paradigms such as Visual Imagery (VI) and Imagined Speech (IS), aimed at addressing challenges like inter-subject variability, BCI illiteracy and improving computational decoding performance. In this context, the present study aims to contribute to the ongoing advancement of non-invasive BCI technology by investigating the potential of Visual Imagery (VI) and Imagined Speech (IS) paradigms. Building upon existing research, this thesis endeavoured to unravel the complexities, advantages, and limitations associated with these paradigms, with a focus on addressing BCI illiteracy and utilising a deep learning model to classify the neural signals. In hope to pave the way for more intuitive and efficient BCI systems, through contributing to the current body of work on VI and IS paradigms.

### 1.2 Literature Review

#### 1.2.1 Advancements in Brain-Computer Interfaces

Over the past few decades, Brain-Computer Interfaces (BCIs) have gained considerable prominence within the field of Neurotechnology. Originally developed to address medical and clinical neurological impairments, BCIs have evolved to encompass a broader spectrum of applications, including commercial endeavours aimed at enhancing cognition, work productivity, and gaming performance. Essentially, BCIs are complex systems capable of deciphering neural signals from users, enabling the translation of intent into actionable commands for computers or machines (Panachekel et al., 2021). Among the various modalities for acquiring neurological data, non-invasive BCI systems that harness electroencephalogram (EEG) recordings offer distinct practical, affordability, and ethical advantages. In contrast to alternative techniques such as functional magnetic resonance imaging (fMRI), magnetoencephalography (MEG), and invasive methods like electrocorticography (ECoG) or intracranial electrodes, EEG-based systems pose fewer ethical concerns. Notably, EEG-based BCIs have been employed to empower individuals afflicted by locked-in syndrome (LIS), a condition in which patients retain cognitive abilities while losing voluntary motor control.

The development trajectory of a BCI system typically involves selecting the cognitive task or mental paradigm that users will perform, followed by identifying neural correlates for accurate decoding. Computational methods, particularly Machine Learning (ML) and Deep Learning (DL) models, are instrumental in classifying the neural data. In current BCI research, an array of paradigms is combined with classical ML models for offline data classification – the process of decoding acquired data. Prominent mental paradigms encompass Motor Imagery (MI), Kinaesthesia, Event-Related Potential (ERP), and Steady State Visually Evoked Potentials (SSVEP). Additionally, emerging paradigms like Visual Imagery and Imagined Speech are gaining recognition. Despite its widespread adoption, MI has encountered challenges in training users to perform the required mental tasks, a phenomenon known as BCI illiteracy, which remains a subject of ongoing investigation (Leeuwis et al., 2021).

#### 1.2.2 Electroencephalogram – Non-invasive Brain Computer Interfaces

The discovery of the electroencephalogram (EEG) dates to 1924, credited to Hans Berger, who pioneered the non-invasive recording of electrical activity from the surface of the human brain cortical neurons. The concept of BCIs harnessing EEG signals was introduced by Vidal in 1973. The type of BCI applications can be divided into three categories: invasive, non-invasive, and hybrid, which combines both approaches. Invasive BCI applications involve complex neurosurgical procedures, ethical considerations, stringent medical regulations, and utilise technologies like intracranial microelectrodes for single and multi-unit neuron recordings, and Electrocorticography (ECoG) (Mridha et al., 2021). In contrast, non-invasive BCI techniques hold broader accessibility as they circumvent neurosurgery. This approach offers benefits such as reduced ethical dilemmas, practicality, and cost-effectiveness. However, it does present its unique set of challenges. Within the non-invasive domain, other neuroimaging, and neurophysiology techniques, including Functional Magnetic Resonance Imaging (fMRI), Magnetoencephalography (MEG), Positron Emission Tomography (PET), and Facial Near Infrared Spectroscopy (fNIRS), are utilised to record brain activity.

Among these modalities, EEG emerges as the most prominent choice for non-invasive BCIs due to its excellent temporal resolution, user-friendly nature, safety, and affordability (Zabcikova et al., 2021). This recent review highlighted the predominance of EEG-based BCIs. It stands as a commercially viable option, and future iterations of commercial BCIs are likely to lean toward EEG-based technology for its user-friendliness and accessibility. Despite these merits, EEG-based BCIs encounter challenges stemming from the methodology itself. Electrodes affixed to the scalp need to capture the collective activity of cortical neurons through the skull, leading to limitations in spatial resolution and signal-to-noise ratio (SNR). Electrode placement requires conductive mediums like gel or saline to improve skin-electrode contact impedance, which can result in signal degradation over time, detachment, and susceptibility to external electrical noise sources like 50 Hz interference, cable movements, and biological artifacts such as eye, muscle, and body movements.

A comprehensive review conducted by Zabcikova et al. (2021) examined 100 original articles concerning current advances and trends in BCI research. Their findings emphasised the prevailing focus of BCI efforts on medical and healthcare applications employing non-invasive EEG-based BCIs. The review highlighted the numerous ongoing challenges and limitations facing BCI systems. These include enhancing signal quality, reducing user training time and effort, developing more comfortable and practical sensors, mitigating muscle and electrical artifacts, enhancing public awareness, streamlining calibration phases, minimising electrode counts, expanding real-time applications, deepening our understanding of the brain, improving classification accuracy, and addressing privacy and ethical concerns. These insights build upon Arico et al.’s (2018) earlier review and underscore the dynamic landscape of BCI research.

#### 1.2.3 Visual Imagery BCI Paradigm

Visual Imagery (VI) is a developing mental paradigm in BCI applications, requiring individuals to conjure mental images of objects, scenes, or individuals. These imagined visuals give rise to neural correlations that can be effectively classified by machines (Nieles et al., 2018). One notable example is Liquidweb’s BCI system, employing the VI paradigm to empower individuals with Locked-In Syndrome (LIS) to communicate with the world. VI involves visualising specific actions, leading to event-related suppression of alpha and beta wave activity. Recent investigations suggest that VI might offer an improved paradigm compared to Motor Imagery (MI), potentially mitigating BCI illiteracy.

In a past study by a Liquidweb student, VI was paired with Echo State Networks (ESN) for classification. The research identified an optimal time window, approximately 1 second after the VI cue, which, when utilised with ESN, enhanced classification accuracy by 20%. However, challenges surfaced, particularly when novice users were tasked with visualising actions in a three-dimensional space, such as pushing a circle into the distance to reduce its size during the ‘push’ cue. This observation underscores the varying levels of difficulty associated with the mental task across participants.

Nieles et al.’s (2018) study aimed to characterise EEG signal patterns during VI of basic shapes like squares, triangles, and circles for potential BCI typing applications. Applying diverse machine learning methods, the researchers intriguingly found no significant correlation between shape types. However, Support Vector Machine (SVM) emerged as the top-performing classifier in terms of accuracy. Their findings suggest that SVM-based classification could serve as a foundation for developing a BCI typing interface using VI for characters and letters.

#### 1.2.4 Imagined Speech BCI Paradigm

Imagined Speech (IS) as a BCI mental paradigm is where the user performs speech in their mind without physical articulation (Panachekel et al., 2021). Previous studies on IS have focussed on types of words used, types of vowels (Tamm et al., 2020), length of words, meaning of words as the stimuli (Lee et al., 2020). Changes in high gamma waves (70-150 Hz) have been identified during overt and covert speech. Overt speech showed high gamma oscillations in the temporal lobe, Broca’s area, Wernicke’s area, premotor cortex, and primary motor cortex. Whereas covert speech showed high gamma changes in the supramarginal gyrus and superior temporal lobe (Pei et al., 2011, Lee et al., 2020, Panachekel et al., 2021, Lopez-Bernal et al., 2022).

Recent strides in the field, such as Lee et al.’s (2020) work, have proposed the symbiotic utilisation of VI and IS for enhanced decoding performance, addressing BCI illiteracy, and practical applications. IS involves users mentally enacting speech without actual articulatory movements. Remarkably, these studies uncovered robust decoding performances for IS and VI, showcasing heightened high-frequency activity in speech and language cortical areas. Classification, facilitated by Support Vector Machine (SVM), indicated that left frontotemporal regions outperformed motor areas for both IS and VI, even in complex 13-class scenarios. As class complexity increased, SVM’s accuracy demonstrated a decrease, prompting suggestions for the incorporation of Deep Learning (DL) models to handle a higher number of classes.

Moreover, Lee et al.’s findings indicated a noteworthy inter participant disparity, highlighting distinct groups excelling in either IS or VI. This variation was postulated to be influenced by unique brain structures or connectivity within individuals. The researchers underscored the pivotal role of selecting an appropriate mental task for successful BCI outcomes. However, they did not delve into strategies to attain this goal, particularly for novice users. This underscores the ongoing quest for refining and personalising BCI experiences.

#### 1.2.5 Addressing BCI Illiteracy

One of the formidable challenges in the realm of BCI applications is the phenomenon known as BCI illiteracy. This intricate issue stems from the difficulty novice users face when attempting to execute designated mental tasks or the extensive training they must undergo, which can become taxing for long-term users. BCI illiteracy is characterized by a user’s inability to effectively carry out the BCI mental task, thereby failing to elicit the anticipated neural correlate change (Leeuwis et al., 2021). This phenomenon has notably challenged well-established BCI mental paradigms like Motor Imagery (MI), kinaesthesia, and Steady State Visually Evoked Potentials (SSVEP).

To combat this challenge, numerous studies have delved into potential factors that might influence users’ aptitude for performing MI-BCI tasks. For instance, a study conducted by Leeuwis et al., (2021) explored the impact of demographic, cognitive, behavioural, and psychological traits on MI-BCI performance. Their findings shed light on key factors that significantly affected performance. Notably, scores on the Vividness of Visual Imagery Questionnaire (VVIQ) emerged as an important indicator. Users who scored higher on the VVIQ, reflecting greater visual imagery characteristics, showcased improved performance in MI-BCI tasks. Furthermore, users with personality traits such as orderliness and autonomy, or a diminished inclination toward fantasy, demonstrated poorer performance on the initial run of MI-BCI. Importantly, these attributes were identified as significant predictors of MI-BCI performance.

The implications of these findings suggest a promising direction for enhancing user specific BCI experiences. By incorporating questionnaires like the VVIQ to gauge user traits, it might become possible to predict which novice users are better suited for specific BCI paradigms, such as Visual Imagery (VI) and Imagined Speech (IS) paradigms. Notably, to the best of the author’s knowledge, the VVIQ has not yet been employed to assess participants’ performance in the VI paradigm. In the current study, the inclusion of the VVIQ aims to evaluate users’ predisposition for visualisation and its potential impact on their VI performance. This endeavour holds promise in potentially mitigating the challenge of BCI illiteracy, allowing novice users to train on mental paradigms that align more closely with their individual abilities.

#### 1.2.6 Classification of EEG signals

For a computer to identify human intent expressed from recorded EEG signals, classification is required. Classification is a supervised machine learning method where the model attempts to predict the correct class of a given input data. The model is first trained using the training data to generalise the data, then trained on test data to validate its classification performance, then fed new unseen data to see how well it can predict the classes.

#### 1.2.7 Machine Learning Models

For the classification of BCI systems, predominantly studies have utilised traditional ML models for classification. The traditional ML models used were mentioned above, however they require long and complex pre-processing and preparation of the data before analysis, are great for binary classification but decline in performance with increased multi-class inputs.

#### 1.2.8 Deep Learning Models

Deep learning (DL) is defined as a combination of machine learning techniques based on artificial neural networks organised in layers (Goodfellow et al., 2016). Recently increased in research and development, it has shown a lot of promise for classification of EEG signals, with benefits in terms of accuracy, multi-class comparisons and reduction of strenuous pre-processing times compared to traditional ML methods. A comprehensive review on decoding IS from EEG-BCI applications by Panachakel et al., (2021), have provided valuable insights to optimising future IS-BCI systems. They concluded that traditional ML techniques only extract features from single channels independently, whereas DL techniques can extract features from channel cross-covariance (CCV) matrices, which can better capture the information transfer between different brain regions. Convolutional neural networks (CNN) are multi-layered neural networks based on the organisation of the cerebral visual cortex (Alzubaidi et al., 2021) and are subdivided into two main parts: feature extraction and classification. The feature extraction layers consist of multiple convolutional layers, reduces the model parameters through convolution and reduces dimension size through pooling. The output of the feature extraction layers produces an extracted feature map that is passed through the fully connected layers for classification. Previous attempts utilising CNN to classify visual objects was 60% accuracy (Llorella et al., 2020). Castro et al., (2020) had success comparing traditional ML classifiers with DL classifiers for VI using the data shapes dataset from Nieles et al., (2018). They found that DL performed better than ML, specifically CNN-Long Short-Term-Memory at 89.44% vs SVM classifier at 83.06%. Indicating DL’s outperformance was through adapting its feature selection and weighting of features to different trials, subjects and tasks which can be translated to IS recognition such as imagine phonemes, imagined words, phrases and sentences.

Mattioli et al., (2021) also had success in implementing a novel DL model, a one-dimensional convolutional neural network (1D-CNN) which achieved high accuracy and transfer learning to classify five classes in MI-BCI. Their approach to transfer learning allowed the 1D-CNN to learn general features from a large population of individuals and was applied to target individuals through a small timeframe of additional training. Based on the success of this model for MI EEG data, this model will be adapted and modified to classify the data collected for VI and IS in this study. Mattioli et al., tested three hyperparameters, Adam optimisation, the use of early stopping, and checkpointing, which was effective to reduce the number of epochs needed for training the system and to reduce the risk of overfitting. Compared to other works (Lun et al., 2020). using the same dataset, their study showed the number of epochs needed for convergence was lower. Batch normalisation (BN) was used in the first two convolutional layers during training and found that without BN the training saw a slower increase in accuracy, loss and was less stable. For data augmentation, synthetic minority over-sampling technique (SMOTE) was used to balance their minority class with the majority class.

### 1.3 Rationale and Objectives

This study will take a comprehensive approach and build on the current foundation of knowledge on VI combined with IS to optimise the decoding performance for BCI use and classification by utilising an adapted 1D-CNN. Moreover, to provide insights into BCI illiteracy and optimal performance in novice users it was explored if there are any correlations of cognitive, behavioural, or psychological factors that can predict VI BCI performance by using the VVIQ.

### 1.4 Hypotheses

Hypothesis 1: During the VI task of visualisation of the “push” condition, alpha and beta suppression would be observed in comparison to the “relax” condition. During the IS task of covert speech of the words “push” and “relax”, gamma oscillations in the temporal and/or dominant speech and language areas of Broca and Wernicke’s would be observed.

Hypothesis 2: Those that score higher on VVIQ will show more affinity towards vividness and imaginary thinking and so will perform better on the VI paradigm compared to those who score lower on VVIQ in novice BCI users.

Hypothesis 3: The 1D-CNN performance would be able to be replicated as in Mattioli et al., for the VI and IS data. Based on the parameters used in that study, such as checkpointing, early stopping and data augmentation, use of the adapted 1D-CNN on this dataset would reproduce similar performance. Furthermore, if there is better performance in one paradigm over the other. This will be measured by the cluster-based permutation statistical analysis and through the CNN’s ability to classify accurately the different neural changes between conditions in VI and IS paradigms.

## 2. Materials and Methods

### 2.1 Participants

Twenty healthy subjects were recruited voluntarily for the experiment (*M_age_* = 27.45 years, SD*_age_* = 2.96 years, 50% female, 5 left-handed) in the local area of London, United Kingdom through word of mouth, online posts, and flyers. The selection criteria included being older than 18 years, having no known neurological conditions such as speech, language and/or auditory impairments, good proficient use of the English language, and were a novice BCI user. Fifteen subjects were right-handed and five were left-handed. The participants were compensated for their participation of 6 GBP. The study was conducted in accordance with the Declaration of Helsinki and the General Data Protection Regulation (GDPR). The experiment protocol was approved by the Research Ethics Committee at Goldsmiths, University of London and in compliance with the Universities UK Research Integrity Concordat. All participants signed an informed consent form.

### 2.2 Materials

#### 2.2.1 Pre-screening Questionnaires

Participants were asked to fill out an online questionnaire (see Supplementary material) containing demographic information (gender, age, dominant handedness, proficiency in the English language, any neurological deficits), Dyslexia questionnaire (Everatt et al., 2002) and Vividness of Visual Imagery Questionnaire (VVIQ) (Marks., 1973) before attending the experiment. The Adult Dyslexia assessment provided from the British Dyslexia foundation was to control for likeability of Dyslexia for the IS paradigm. It contains 15 questions in total, where participants rate how true each statement is to them. The overall score gives an indication of likelihood of Dyslexia symptoms; scores less than 45 were “probably non-dyslexic”. The VVIQ measured vividness of visual imagery used in Leeuwis et al., (2021) for the VI paradigm. It quantified the intensity to which people can visualise settings, persons, and objects in mind. It contained 16 questions, of which the mental image for each was rated along a 5-point Likert scale, where 5 indicated “perfectly clear and vivid as normal vision,” and 1 indicated “no image at all.” The item scoring was as per Leeuwis et al., where a higher VVIQ score (maximum 80) indicated a higher vividness of the mental image.

#### 2.2.2 Data Acquisition

The EEG data was collected using a 64-channel BioSemi ActiveTwo, with a sampling rate of 1024 Hz. Following the international 10-20 system, the electrodes were placed at Fp1, AF7, AF3, F1, F3, F5, F7, FT7, FC5, FC3, FC1, C1, C3, C5, T7, TP7, CP5, CP3, CP1, P1, P3, P5, P7, P9, PO7, PO3, O1, Iz, Oz, POz, Pz, CPz, Fpz, Fp2, AF8, AF4, AFz, Fz, F2, F4, F6, F8, FT8, FC6, FC4, FC2, FCz, Cz, C2, C4, C6, T8, TP8, CP6, CP4, CP2, P2, P4, P6, P8, P10, PO8, PO4, O2 with Common Mode Sense (CMS) active electrode and Driven Right Leg (DRL) passive electrode as the ground electrodes. The device used sintered Ag-AgCl electrode tips, placed into a headcap where the electrode ports were filled with conductance gel. Four external electrodes were placed, two behind bilateral mastoids for the reference and two for the heart rate: below the right shoulder and above the left hip bone. The recording parameters were low-pass: 150 Hz and high-pass: 0.1 Hz. Impedance varied across participants but was kept below 40 kOhms overall. The computer screen was put on night vision brightness to reduce eye strain. Troubleshooting to reduce impedance and artefacts, included re-gelling the electrodes, checking electrodes were placed well, and replacing electrodes sets.

#### 2.2.3 Stimuli and tasks

In collaboration with LiquidWeb, the data collection protocol was adapted from their existing Visual Imagery (VI) paradigm, with addition of the Imagined Speech (IS) paradigm. This was performed through Python coding language and the PsychoPy module in VS Code.

LiquidWeb’s standard data protocol: “Push” and “Relax” were presented as visual cues of the words “Push” and “Relax” in black font against a white background (Figure 1). The choice of visual prompt was kept the same between the two paradigms, the defining difference between the two paradigms was the different mental tasks performed. This was explained through the instructions (Figure 1). PsychoPy displayed the VI and IS prompts on a computer screen in a dark sound proofed room. The instructions of both paradigms were explained to the participants before the experiment, and they were shown a video for the ‘Push’ condition in VI (Figure 1).

**Figure 1.**
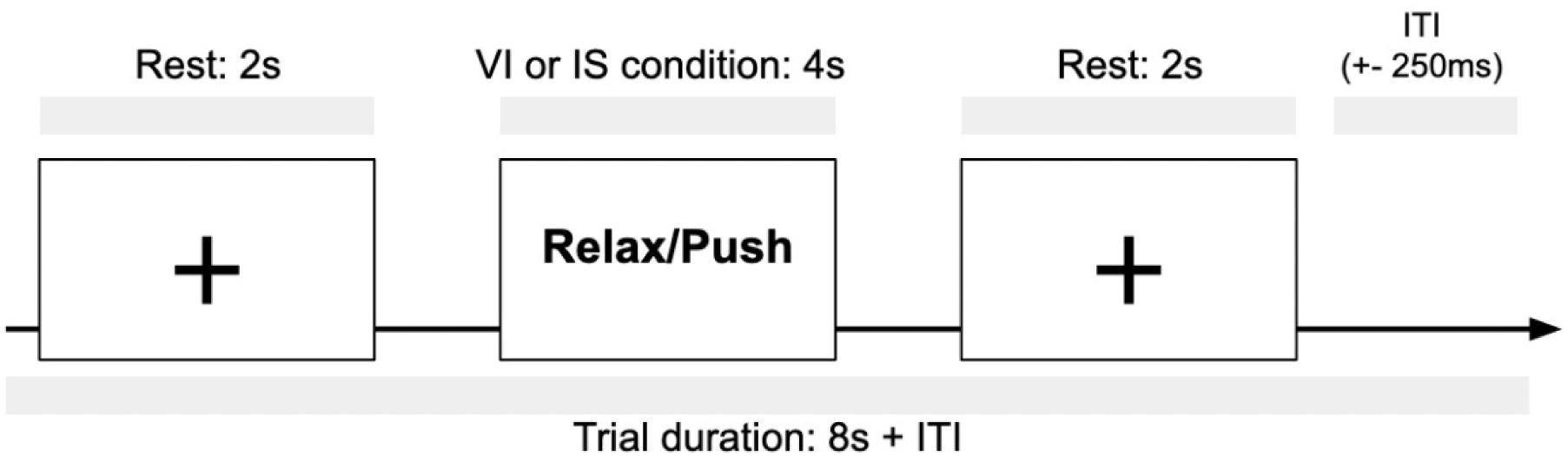
Diagrammatic representation of a trial in the data collection protocol. The VI block instructions: A fixation cross indicated a neutral state, blinking allowed. During ‘Relax’ condition: Participants were asked to imagine a scenery, object or feeling of relaxing. During the ‘Push’ condition: Participants were asked to imagine pushing an object into space, making it smaller. The IS block instructions: A fixation cross indicated a neutral state, blinking allowed. During ‘Relax’ condition: participants were asked to imagine speaking the word shown on screen in their mind, without physically articulating. Repeating approximately once every second. During the ‘Push’ condition: participants were asked to imagine speaking the word shown on screen in their mind, without physically articulating. Repeating approximately once every second. Each block consists of 50 trials, 25 trials in the ‘Push’ condition and 25 trials in the ‘Relax’ condition, the order of condition was randomised. Each participant would have completed 100 trials in the VI paradigm and 100 trials in the IS paradigm, totalling 200 trials. The participants were instructed to keep as still as possible during the experiment, reduce blinking as much as possible when the prompts were presented, and to blink when the fixation cross was shown. In each trial, the fixation cross was shown 2-seconds preceding the 4-seconds prompt, and 2-seconds after. After 25 trials, the fixation cross was shown for 15 seconds, which allowed the participants a break.

**Figure 2.**
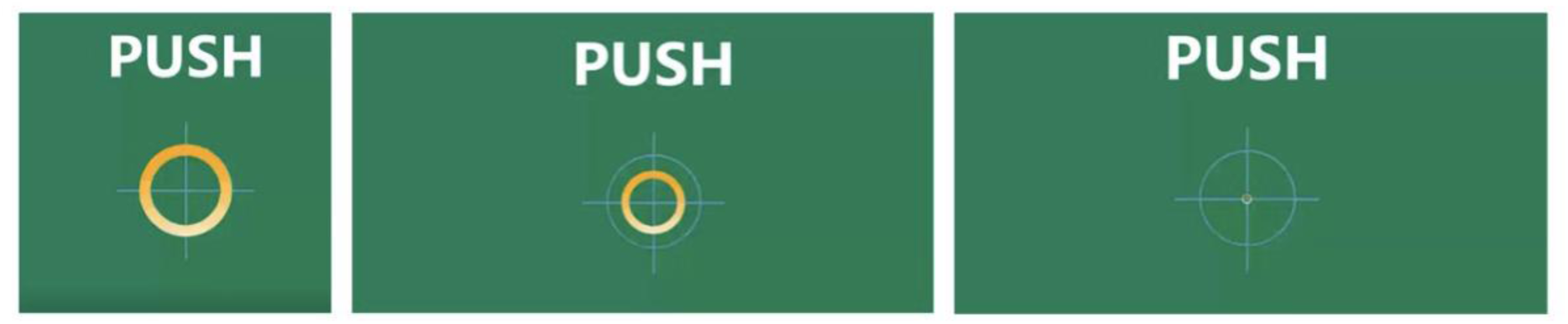
Screenshots of the Visual Imagery ‘Push’ condition instruction video. The video shows a visual representation of the “push” task, where the user can picture in their mind pushing the circle into the distance so that it becomes smaller. This was displayed on a computer to show the subjects prior to the experiment.

### 2.3 Experimental Procedure

The participants were sent the online questionnaire and online consent form prior to attending the experiment. They were asked to refrain from drinking caffeine and alcohol 24 hours prior to attendance. On the experiment day, participants had the EEG headcap set up and then they went to a noise isolated dark room to perform the data protocol tasks. Figure 1 outlines the experiment procedure. The data protocol consisted of five blocks in total: one 5-minute baseline, two consecutive blocks of VI and two consecutive blocks of IS, the paradigm order counterbalanced manually between participants to account for interference of one paradigm over the other. It was not feasible to separate the two paradigms into two sessions as in Lee et al., (2020) due to limited availability of the EEG lab and time constraints. The participants were given breaks after each block of a few minutes, enabling them to move, stretch, have some water, and give feedback to the experimenter. The task instructions were reiterated to the participants before each block and during their break. At the end of the experiment, the EEG headcap was removed, and the participants were debriefed.

### 2.4 Data Pre-processing

The pre-processing pipeline was programmed using MNE-Python, an open-source Python library for EEG and MEG analyses (Gramfort et al., 2013), following standard EEG pre-processing guidelines. The EEG data was recorded at 1024 Hz, it was down sampled to 256 Hz prior to analyses. The continuous raw EEG data was filtered to improve signal to noise ratio (SNR) with 50 Hz notch filter, 0.5 Hz high-pass filter and 125 Hz low-pass filter to capture gamma band oscillations (Lee at al., 2020). The raw EEG data was then visually inspected for each participant, bad channels that showed drifting, sweat artefact, and movement artefact were interpolated. The data was then separated into relevant epochs. The data was time-locked to the onset of the cue of interest (−1, +4s). The epochs were visually inspected and noisy epochs were rejected. Artefact correction was performed with Independent Component Analysis (ICA) to remove the typical EEG biological artefacts such as eye blinks and muscle artefacts. The extracted epochs arrays were nEpochs*nChannels*nTime.

### 2.5 Data Analysis

#### 2.5.1 Questionnaire Statistics

The online questionnaire was separated out into VVIQ and Dyslexia scores using R Studio. The Dyslexia questionnaire results were calculated for each participant and compared with the scale of likeability for Dyslexia. The VVIQ questionnaire results for each participant was totalled, the group mean score was calculated and the participants were divided into two groups: High VVIQ group, those that scored above the mean score and Low VVIQ group, those that scored below the mean score. 14 participants were in the High VVIQ group, and 6 participants were in the Low VVIQ group.

#### 2.5.2 Time Frequency Analysis – Morlet Wavelet Convolution

For feature extraction of high dimensional EEG data, time-frequency (TF) analysis was performed. TF analysis is a relatively new method in the field of cognitive neuroscience, which has shown ability to characterise the temporal dynamics of the features in oscillations contained in EEG data: frequency, power, and phase. TF measures can provide a closer interpretation of neurophysiological mechanisms by separating power and phase information across different frequencies, as opposed to event related potentials (ERPs) and Fourier-based analyses (Morales & Bowers., 2022). The TF decomposition in both VI and IS paradigms was performed with MNE Python. The pre-processed epochs were separated into the four conditions: VI Relax, VI Push, IS Relax and IS Push. The TF components were extracted using Morlet wavelet convolution, at 5 cycles and 8-125 Hz to consider the three band frequencies of interest, alpha, beta and gamma. Baseline correction was applied between –0.5 and 0.0. The TF components were averaged across alpha (8-12), beta (12-30) and gamma (30-125). Then averaged across trials to give *nC*ℎ*annels* ∗ *n*Fr*equency* ∗ *nTime*. The arrays per subjects were then concatenated per the four conditions. The result was four arrays of *nSubjects* ∗ *nC*ℎ*annels* ∗ *n*Fr*equency* ∗ *nTime* to extract power as a function of time and space. The result was plotted for alpha, beta and gamma for the four conditions. The extracted TF decomposition was then statistically analysed with non-parametric cluster-based permutation analysis. The arrays were saved into NumPy format for the non-parametric cluster-based permutation analysis.

#### 2.5.3 Non-parametric Cluster-based Permutation Analysis

Non-parametric cluster-based permutation analysis have been widely used in current cognitive neuroscience practices to provide statistical analysis for the complexity of high-dimensional data from EEG (Sassenhagen & Drashkow, 2018). It was first proposed by Maris and Oostenveld (2007), to address the multiple comparison problem observed with utilising parametric statistical methods for EEG data. In addition, cluster-based permutation analysis delivers high Type II (power) and has been effective in controlling for nominal Type I (false positive) error rates for the null hypothesis of exchangeability considering the cluster structure of the data. To compare the neural frequency oscillation changes between the conditions ‘relax’ and ‘push’ in both VI and IS paradigms, non-parametric cluster-based permutation tests (two-sided with an α level of .025 (Maris, & Oostenvald., 2007). To perform the paired permutation analyses for within-group comparison, the four TF NumPy arrays were converted into MATLAB files, to be analysed using FieldTrip, in MATLAB_R2022b (1999). The analysis was performed within paradigms e.g., VI Relax vs VI Push and IS Relax vs IS Push. Each frequency band, alpha, beta and gamma were analysed. The parameters used for the analyses were the Monte Carlo method, 1000 permutations, significant level of p=0.025 and the latency window at 1.0-3.0 seconds; chosen based on the alpha and beta suppression shown from the TF analysis.

### 2.6. 1D-CNN model for Classification

#### 2.6.1 One-dimensional convolutional neural network structure (1D-CNN) and function

For classification of the VI and IS paradigms for BCI applications, the 1D-CNN computational model was adapted from Mattioli et al. (2021) mentioned above. The 1D-CNN model was implemented using the Python TensorFlow module. The code scripts of the model are included in supplementary materials. 1D-CNN means during convolution the CNN kernels slide only over the elements of 1 dimension of the input pattern; time. The model takes the input *M* × N where *M* is the length of the time window considered and N is the number of EEG channels (Mattioli et al., 2021). Each 1D convolutional layer uses a kernel of variable dimension input *Q* × N where *Q* is the temporal window covered by the filter and N is the number of channels. The mathematical notation of a 1D convolutional layer is:

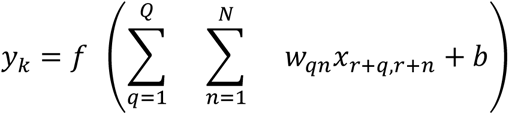

(1) where *y*_r_ is the output of the unit r of the filter feature map of size R (R = *M* in the case stride=1 and ‘padding’ is used), *x* is the two-dimensional input portion overlapping to the filter; *w* is the connection weight of the convolutional filter; *b* is the bias term and *f* the activation function of the filter. The dimension of the filter feature map after the convolution operation (R) was calculated by:

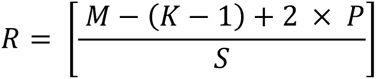

(2) where P is the padding size; *S* is the stride.

The overall features of the adapted CNN architecture are shown in Figure 3. Please refer to Mattioli et al., (2021) for in depth explanation of mathematical functions for each layer. To summarise, the adapted 1D-CNN used for this study, used different dimensioned input matrix for the VI and IS paradigms, due to the different datasets. In addition, the activation function for the final layer was changed to sigmoid to suit the binary classification within VI and IS paradigms.

**Figure 3.**
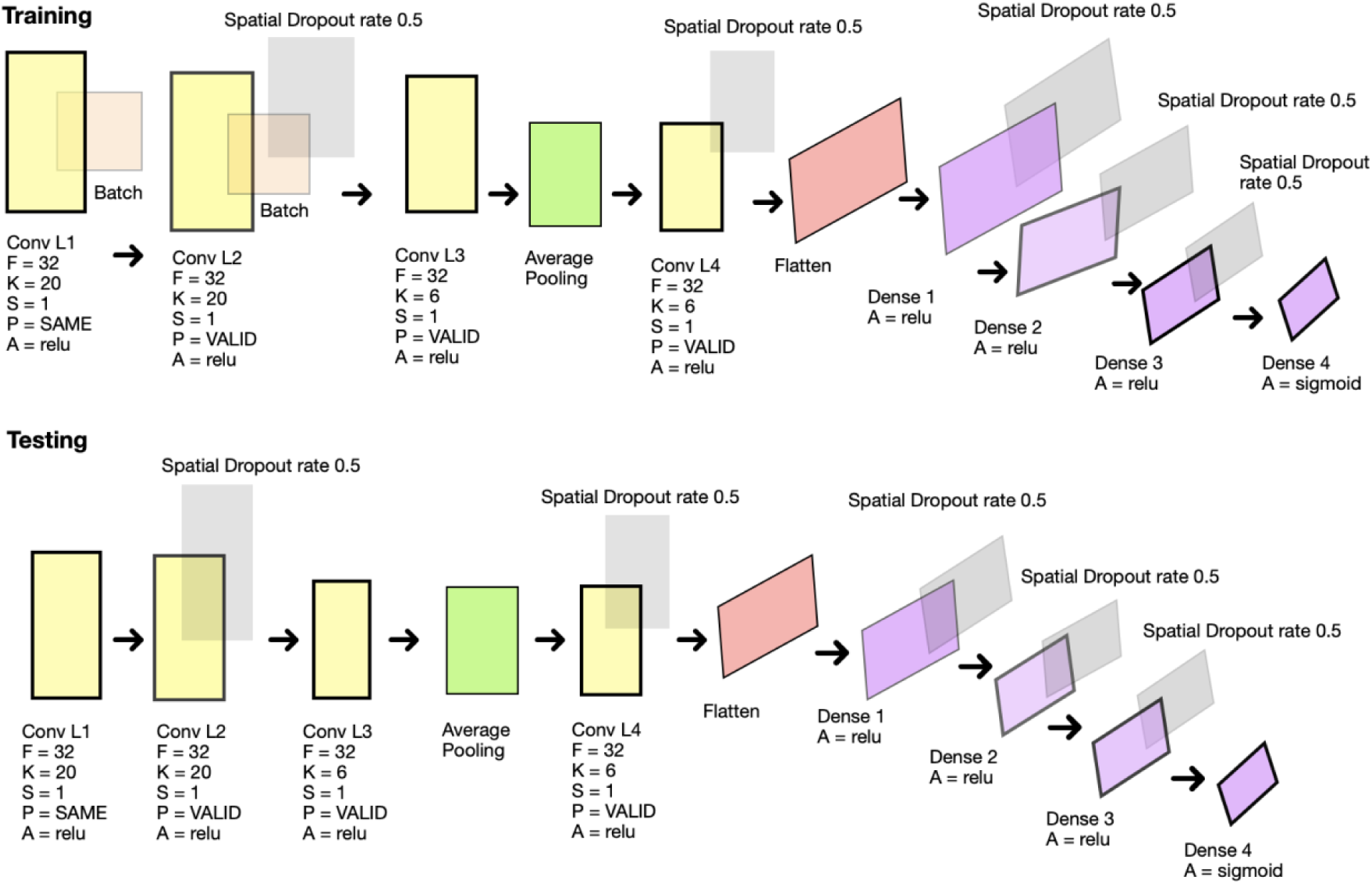
1D-CNN architecture diagrammatic representation. This is the adapted 1D-CNN model used for classification of EEG-based BCI for VI and IS paradigms. Layers 1-10 from the left. Training utilised batch normalisation whereas testing did not. Conv L1-L4 = Convolutional layers 1-4, Batch= batch normalisation, F= filters, K= filter size, S= stride, P= padding (TensorFlow definitions), A= activation function, Average Pooling= 2 × 1 with stride of 2, Spatial dropout rate= 0.5 (p=50%), Flatten= reshapes the matrix to vector, Dense 1-4= fully connected units.

Figure 3 summarises the 1D-CNN architecture. There were ten layers, the first convolution layer (L1) kept the exact size of input and output. Batch normalisation (BN) is applied after L1, to normalise and zero-centre the entire batch input, allowing the model to learn the optimal scaling of the input. Layer 2 (L2) is the same as L1, but without padding applied. BN is applied again and then spatial dropout (SD) for additional regularisation. This was achieved during the training phase by excluding units with probably p=50% at each training step. Layer 3 (L3) performs convolution again with a smaller kernel. Followed by 1D average pooling of the input, by 2 × 1 to reduce the size of each dimension by a factor of 2. To reduce the computation and network parameters needed. Layer 5 (L5) is the 4^th^ convolution with SD. Layer 6 (L6) is the flatten layer, where the matrix input is reshaped to a vector for processing in fully connected layers. Layers 7, 8 and 9 were fully connected with SD, rectified linear unit (ReLU) activation functions. Layer 10 used a sigmoid function for classifying the two classes within each paradigm (VI and IS).

#### 2.6.2. Dataset and data augmentation

The data was separated out into the two paradigms VI and IS, and binary classification was performed within each paradigm between “relax” and “push” classes. This was because following the non-parametric cluster-based permutation analysis, only significant time-frequency was observed in the VI paradigm and not in the IS paradigm. For the VI paradigm the 1D-CNN was fed the input *M* × N where *M* was the significant time window between 1.5-3.0 seconds multiplied by the sampling rate (SR) 256 Hz, and N was 46 the number of significant channels. For the IS paradigm, there was no significant time-frequency hence the whole dataset input, *M* was 4.0 seconds multiplied by 256 Hz, and N was 64 the number of channels.

Data augmentation was generated to increase the dataset per paradigm, per participant as there were approximately only 100 trials in VI and 100 trials in IS. This was necessary to use the 1D-CNN to help the model generalise better as there was not enough data. Mattioli et al., (2021) used synthetic minority over-sampling technique (SMOTE) but in this study, random cropping training was used instead to avoid overfitting due to limited data. Random cropping training was performed in the temporal domain, where synthetic random data was generated by cropping the time length of the original data (Schirrmeister et al., 2017). The function was defined with the input arguments of x array: *nEpoc*ℎ*s*, *nC*ℎ*annels*, *nTime*, y array: *nEpoc*ℎ*s* label array, SR: 256 Hz, segment length: the length of the new segment in seconds (defaults to 1) and overlap factor: the overlapping factor between segments (defaults to 0). The output was an array of *nAugmentedExamples*, *nAugmentedEpoc*ℎ*s*, *nC*ℎ*annels* where the cropped time increased by the factor of segment length. This was performed on the whole dataset, including the training, validation, and test set.

The dataset was then randomly split into three sub-datasets where 80% of the data formed the training set, 10% the validation set and 10% the test set. The splitting was stratified in a balanced manner, to ensure the model had no knowledge of the data during testing. One hot encoding was then applied to the three sub-datasets to encode the classes to integers of ‘0’ and ‘1’ for the 1D-CNN.

#### 2.6.3 Network Training

The 1D-CNN classification was trained separately for each subject, per paradigm. The optimisation of the 1D-CNN defined the loss function with binary cross-entropy which calculates the loss between the predicted classes and true classes. The Adam optimisation is a stochastic gradient descent (SGD) algorithm based on a different learning rate for each parameter which adapts based on the first order and second order moments of the gradient. It was used to minimise the binary cross-entropy loss and to update the CNN parameters (Kingma & Ba., 2015). As per Mattioli et al., (2021), the hyperparameters of the algorithm set at: α = 0.0001, β_1_ = 0.9, β_2_ = 0.999, ϵ = 10^−8^. Where the error gradient was calculated for batch size of 10. Training epoch range was set at 100, however the following methods defined the resulting number of epochs trained. Validation early stopping was used to avoid overfitting, when the loss on the validation set does not improve for at least a threshold value (0.001) for 10-20 consecutive epochs. Check-pointing technique was used to save the parameters of the 1D-CNN at *e*_*t*_. If the validation loss did not improve in epoch *e*_*t*_, the parameters of the previous epoch *e*_*t*−1_ were used for the next epoch *e*_*t*+1_, discarding epochs of lowering performance. Training and testing were performed on a MacBook computer using the M1 Silicon chip. The training of 1D-CNN for a given subject was approximately 10 minutes.

#### 2.6.4 Performance metrics

The performance of the VI and IS paradigms and for each class “relax” and “push” were measured through four standard indexes: precision, recall, accuracy, and F1-score.

Precision for a given class:

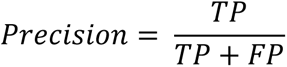

Recall for a given class:

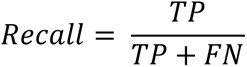

Accuracy for a given class:

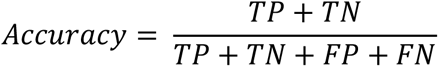

F1-score for a given class (the index is the harmonic average of Recall and Precision):

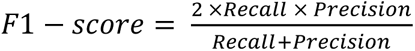

The indexes are defined by the ratios between the frequencies of the cells of the confusion matrix for binary classification: the rows reporting the true classes and the columns reporting the predicting classes. True positive (*T*P) indicate the instances belonging to the positive class being classified correctly. True Negative (*T*N) indicate the instances belonging to the negative class being classified correctly. False Positive (FP) indicate the instances belonging to the negative class but wrongly classified as belonging to the positive class. False Negative (FN) indicate the instances belonging to the positive class but wrongly classified as belonging to the negative class.

## 3. Results

### 3.1 Questionnaire results

The results of the questionnaire showed all but one of the participants scored below the “probably non-Dyslexic” range (0 to 45), with subject 9 on the minimum threshold (48) of “showing signs consistent with mild Dyslexia” range (45 to 60). The mean VVIQ score was 55.6 (SD = 15.31), the highest score was 75 (subject 14) and the lowest score was 16 (subject 2). The groups were split into the high VVIQ group and low VVIQ group by those who scored above the mean score and below the mean score, respectively (Figure 4).

**Figure 4.**
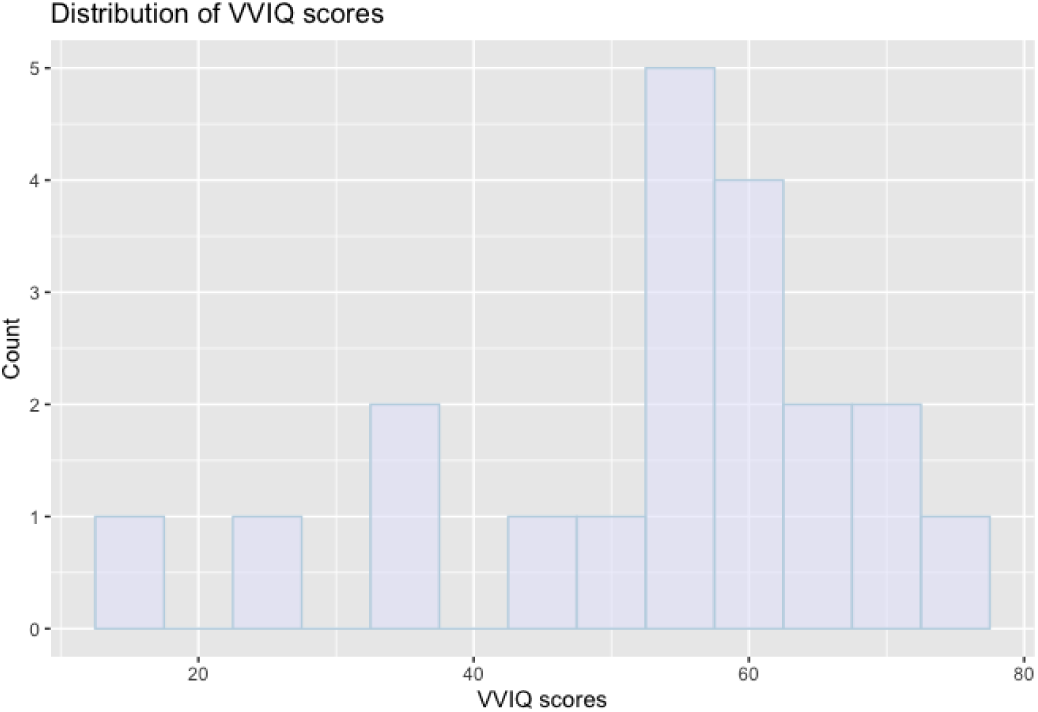
Distribution of VVIQ scores. Histogram showing the distribution of VVIQ scores of all subjects.

### 3.2 Time Frequency and Nonparametric Cluster-based permutation Analysis

To assess significant neural oscillations during the VI paradigm of alpha and beta frequencies, time frequency and non-parametric cluster-based permutation analyses were performed. The VI paradigm showed alpha and beta suppression in the push condition compared to the relax condition, which was statistically significant *t* = 2.035^+04^, *SD* = 0.003, *CI* = 0.005, *p* =0.009. For the alpha frequency band, the significant suppression occurred between 1.61 – 3.0 seconds after the onset of stimuli cue. For the beta frequency band, the suppression occurred between 1.51 – 3.0 seconds after the onset of stimuli cue (Figure 5). The significant channels are highlighted by the topographic maps in Figure 5. The hypothesis that there would be alpha and beta suppression during the VI mental task was supported, and the null hypothesis of no significant difference between the two states was rejected. The same was performed for the IS paradigm for alpha, beta, and gamma frequencies. However, there was no statistically significant change between “push” and “relax” conditions for alpha, beta, or gamma oscillations, as shown in Figure 5.

**Figure 5.**
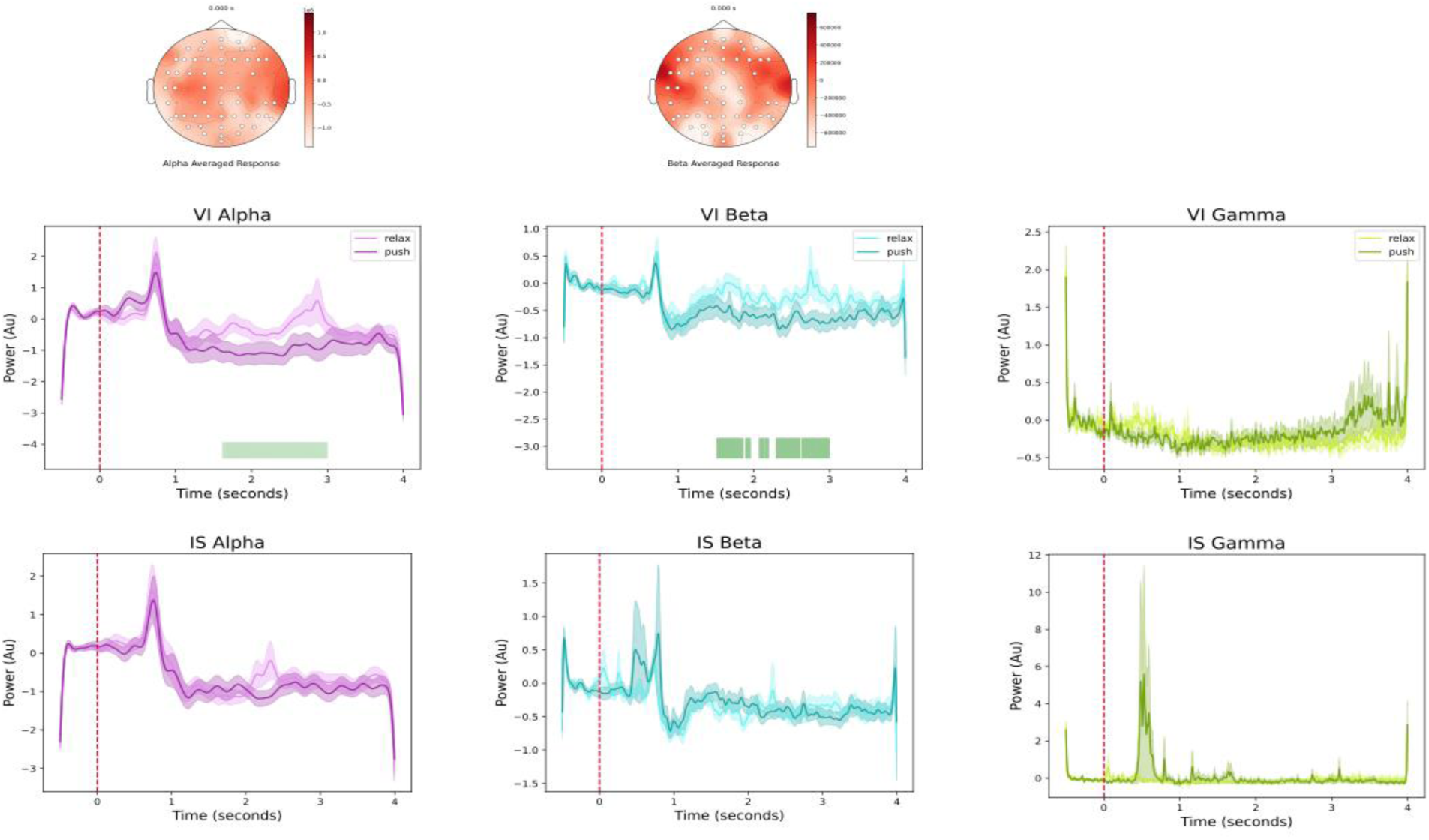
Time-frequency analysis plots of Visual Imagery vs Imagined Speech paradigms. The TF plots were generated by extracting the relevant frequency ranges for alpha, beta, and gamma through Morlet wavelet convolution (see Methods). The onset of the cue was presented at time point ‘0’, marked by the dotted red line. The power of each frequency is in arbitrary units. The topographic map shows the significant channels where alpha and beta suppression was observed during the push condition, shown by the white dots. Black dots show non-significant channels.

#### 3.2.1 High VVIQ group vs low VVIQ group

The hypothesis that those who scored higher on the VVIQ questionnaire would be able to perform the VI paradigm tasks better than those who scored lower, hence performing better on the VI paradigm was supported. This was measured by the significant alpha and beta suppression in the push condition compared to the relax condition in the high VVIQ group. The high VVIQ group showed alpha and beta suppression in the “push” condition compared to the “relax” condition; this was statistically significant, *t*= 2.008^+04^, *SD*= 0.003, *CI*= 0.0059, *p*=0.009. For the alpha frequency band, the significant window occurred between 1.23 to 2.54 seconds and for the beta frequency band, the significant duration window between 1.31 to 2.57 seconds (Figure 6). There were no significant neural changes in all three frequency bands in the IS paradigm for the high VVIQ group (Figure 6). Figure 7 shows the low VVIQ group, where the VI paradigm did not show any significant difference between push or relax conditions in all three frequency oscillations, alpha, beta (*t*= 1.285^+03^, *p*= 0.19) and gamma (*t*= –87.22, *p*= 0.44). This was similar to the IS paradigm, there was no significant difference in all three frequencies alpha, beta (*t*= 552.48, *p*= 0.52) and gamma (*t*= –55.15, *p*= 0.40).

**Figure 6.**
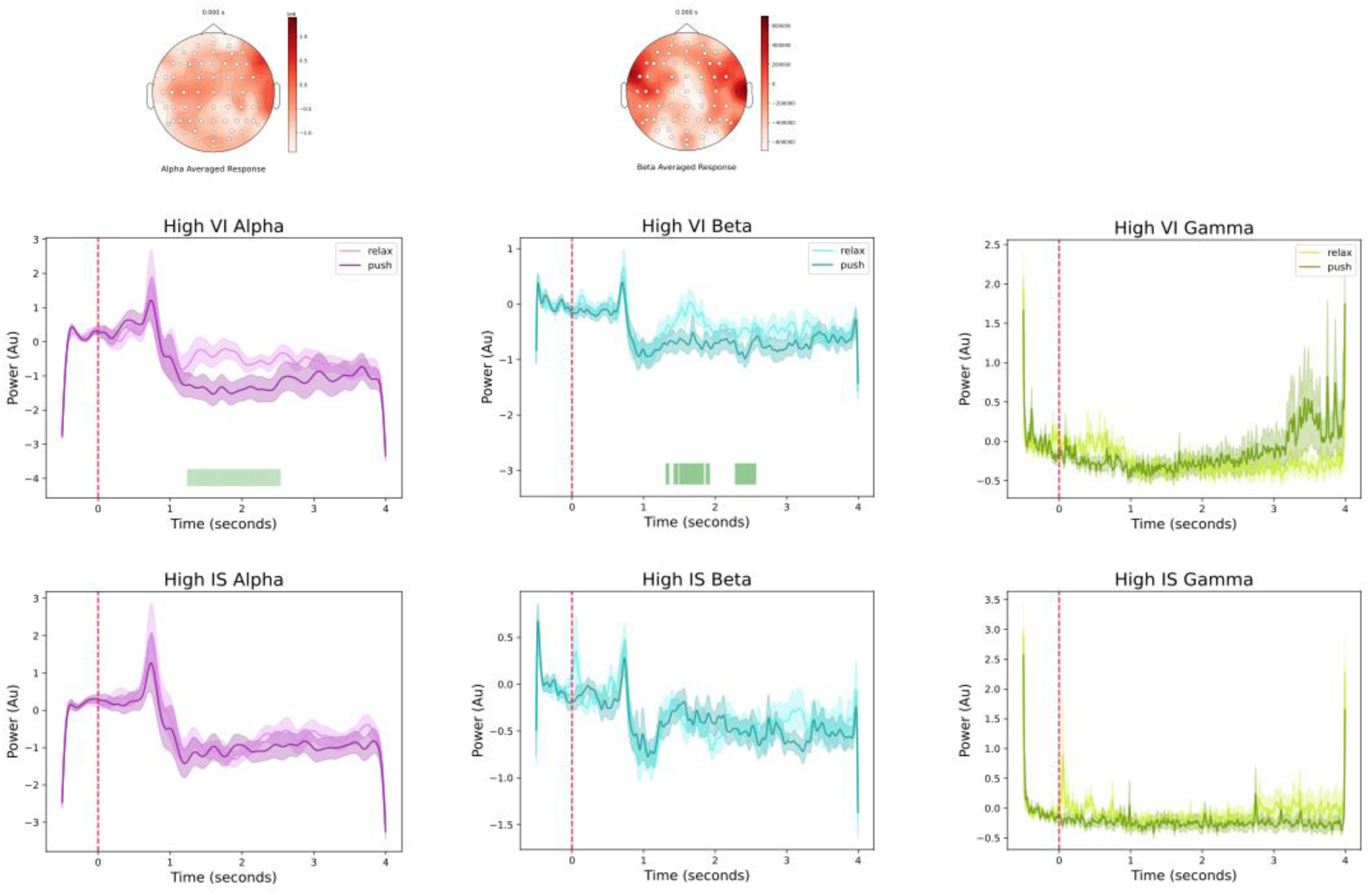
Time-frequency analysis plots of high VVIQ group in VI and IS. The TF analysis was extracted for the high VVIQ group for alpha, beta, and gamma frequencies. The onset of the cue was presented at time point ‘0’, marked by the dotted red line. The power of each frequency is in arbitrary units. The topographic map shows the significant channels where alpha and beta suppression was observed during the push condition, shown by the white dots. Black dots show non-significant channels.

**Figure 7.**
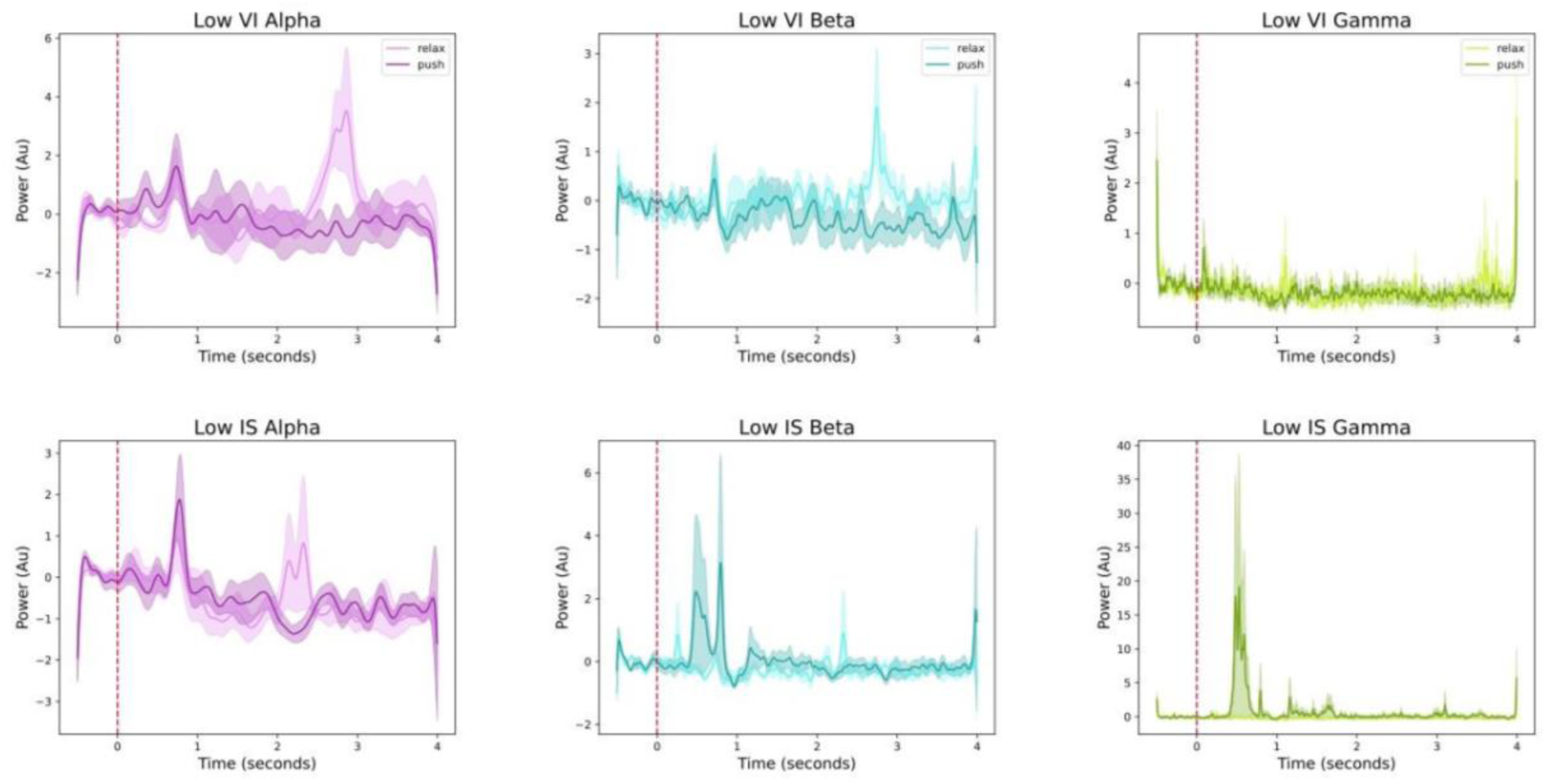
Time-frequency analysis plots of low VVIQ group in VI and IS. The TF analysis was extracted for the high VVIQ group for alpha, beta, and gamma frequencies. The onset of the cue was presented at time point ‘0’, marked by the dotted red line. The power of each frequency is in arbitrary units.

### 3.3 Feature Extraction and Classification

#### 3.3.1. 1D-CNN overall classification performance

The 1D-CNN model was trained by changing the overall factor parameter in the random cropping data augmentation which provided optimal model performance overall. It was observed that increasing the overall factor from 0.5 to 0.9 increased the accuracy of the 1D-CNN from approximately 50% to 99% respectively. It was observed that increasing the patience parameter of the validation early stopping to 20 epochs for lower overlap factors 0.5-0.6 allowed more epochs through to train initially. At an overlap factor of 0.9, a patience of 10 allowed faster convergence, with less epochs required to achieve 90-98% accuracy. After this initial training the training parameters for all datasets were kept consistent at overlap factor of 0.9 and patience of 10. Table 1 shows the results of the classification for the two paradigms VI and IS. The table reports the number of epochs, the test set final loss, and the performance indexes computed on the test set and averaged over all the subjects. The 1D-CNN showed overall 89.3% accuracy in performance for VI and 77.7% accuracy performance for IS, VI showed higher accuracy performance than IS by 11.6%. The performance metrics for the “relax” and “push” class for the VI and IS supported the respective accuracy scores. A paired t-test was then performed between the mean accuracy performance of VI and IS, which was statistically significant, *t*(19) = 2.89, *p* = 0.009.

**Table 1.** Classification report of the 1D-CNN overall performance for Visual Imagery (VI) and Imagined Speech (IS). It includes the training epochs, final test-set loss, average accuracy per paradigm (VI and IS), and performance indexes of the classes (Relax and Push) averaged over all subjects and computed with the test set.

| Metrics\ Paradigm | VI |  | IS |  |
| --- | --- | --- | --- | --- |
| Epochs number | 58.55 |  | 38.45 |  |
| Test-set loss | 0.24115 |  | 0.4106 |  |
| Avg Accuracy (%) | 89.32 |  | 77.695 |  |
|  | **Relax** | **Push** | **Relax** | **Push** |
| Avg precision (%) | 89.75 | 89.7 | 79.5 | 77.65 |
| Avg recall (%) | 88.95 | 89.8 | 74.6 | 80.8 |
| Avg F1-score (%) | 89.05 | 89.45 | 75.8 | 78.65 |

#### 3.3.2. 1D-CNN classification performance with data augmentation

The classification performance of the 1D-CNN was then assessed with the loss and accuracy plots between the best and worst performing subjects in VI and IS. For VI, the best was subject 12 and worst was subject 1 (Figure 8). For IS, the best was subject 4 and worst was subject 19 (Figure 9). The result of these plots was not what is expected for a model of good fit, the plots show that there is overfitting, given by the gap between the validation loss and accuracy compared to the training loss and accuracy, respectively. In addition, the unstable nature indicated uncertainty for the 1D-CNN to predict the validation set. This suggests that there is under representation of the validation set. However, comparison of the best to worst subject performance showed that the worst performing participants in both paradigms also have poor representing plots.

**Figure 8.**
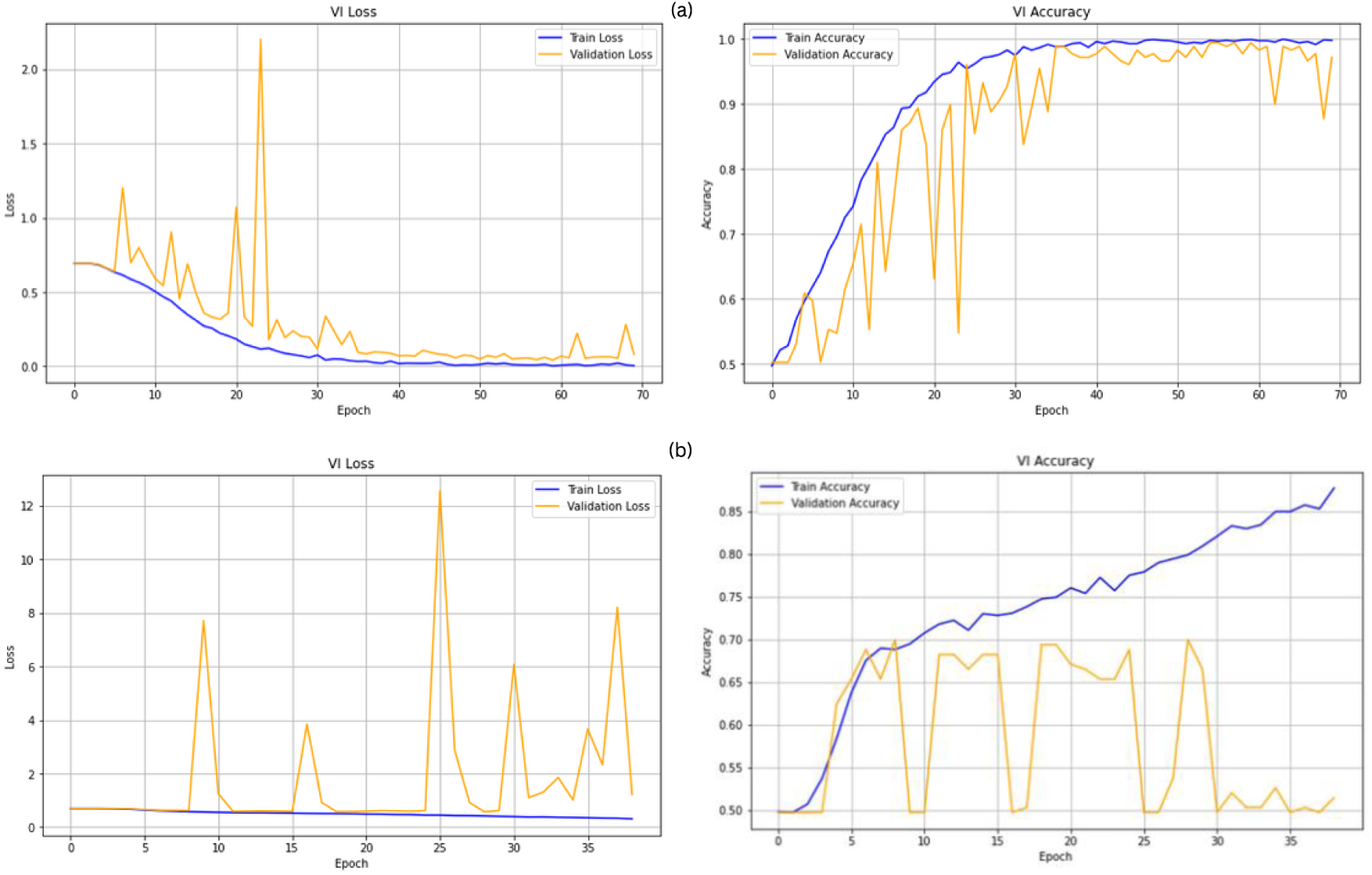
1D-CNN Classification performance of Visual Imagery comparison between best and worst subjects. Loss and accuracy on (a) best performing subject 12 and (b) worst performing subject 1, during the training epochs, computed on the training and validation sets.

**Figure 9.**
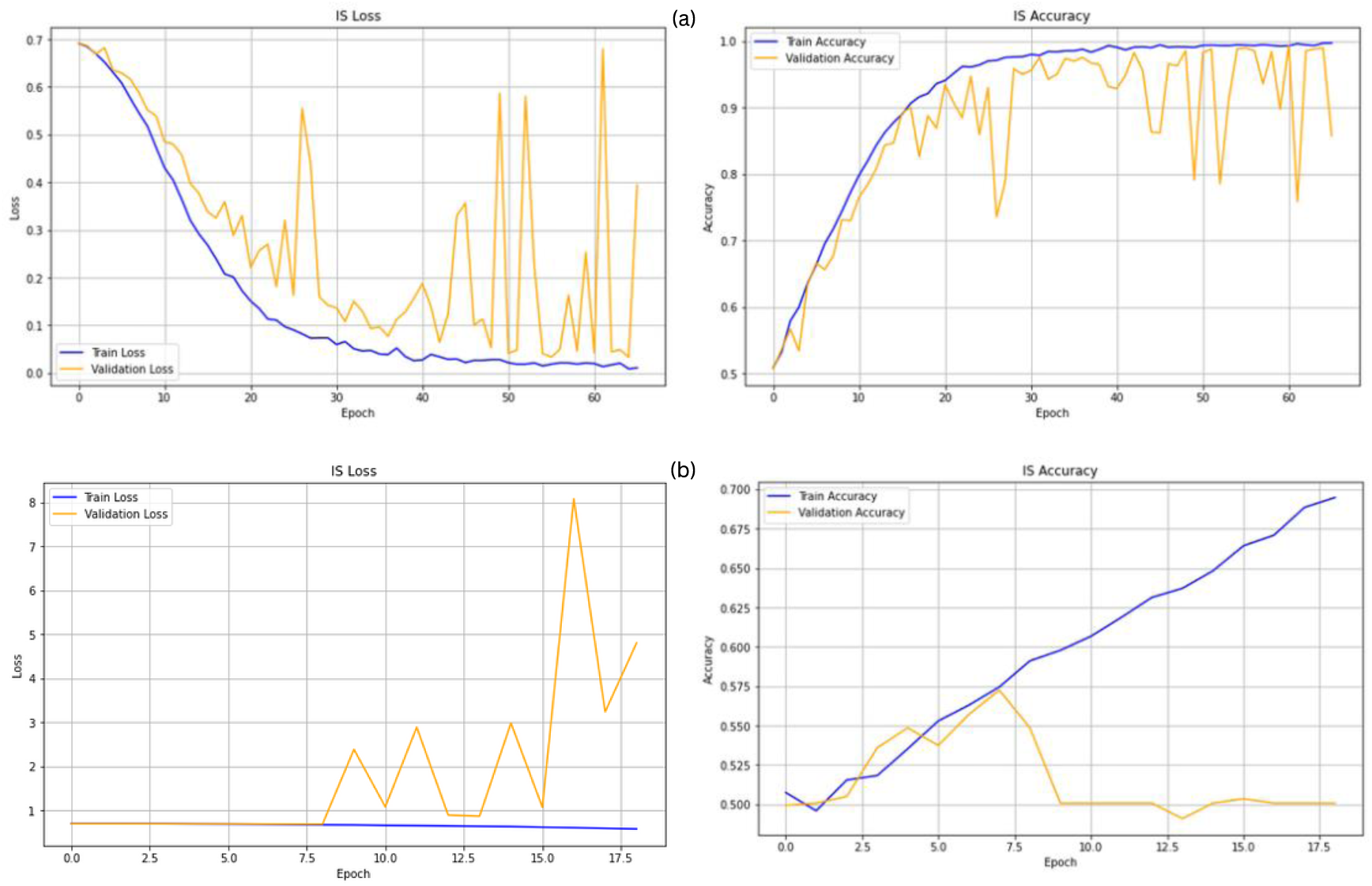
1D-CNN Classification performance of Imagined Speech comparison between best and worst subjects. Loss and accuracy on (a) best performing subject 4 and (b) worst performing subject 19, during the training epochs, computed on the training and validation sets.

#### 3.3.3 VVIQ scores vs VI 1D-CNN Accuracy

To further assess the impact of VVIQ scores, a linear regression model was performed to compare if there was a significant relationship between high VVIQ scores and VI 1D-CNN classification accuracy performance. It showed there was no statistically significant relationship, R^2^ = –0.05, *F*(1, 18) = 0.0015, *p* = 0.96 (Figure 10).

**Figure 10.**
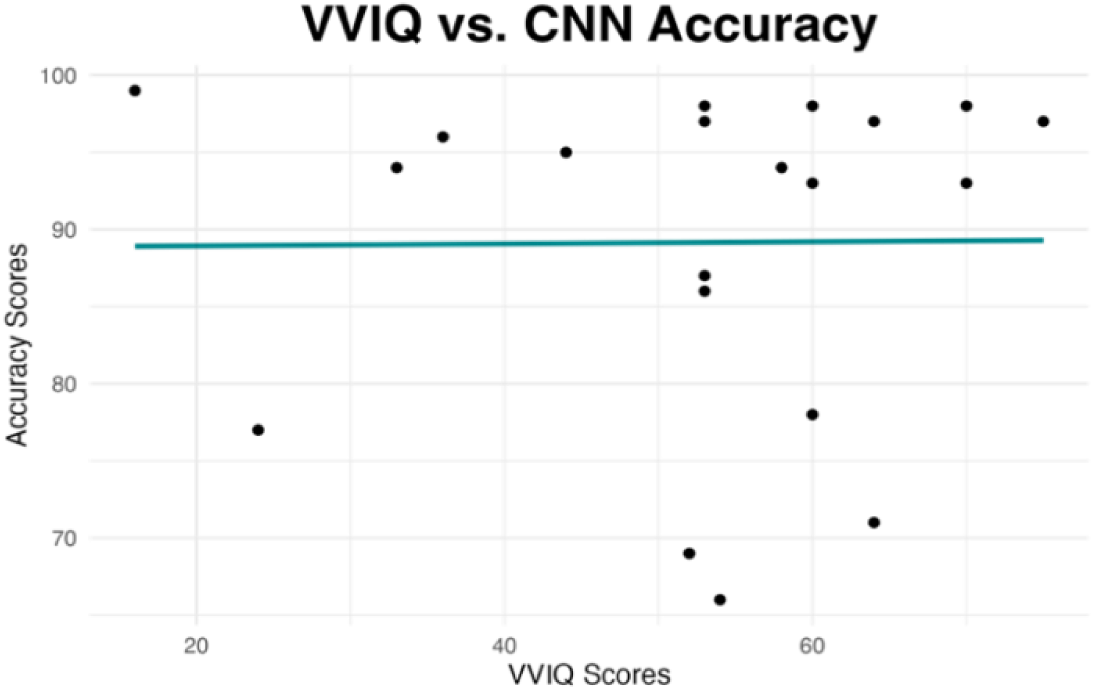
VVIQ vs 1D-CNN Accuracy. A linear regression model was performed to compare the individual VVIQ scores to the 1D-CNN VI accuracy scores for each participant.

## 4. Discussion

### 4.1 Visual Imagery BCI

The hypothesis that the VI paradigm would produce alpha and beta suppression in the “push” condition compared to the “relax” condition was supported, the results showed significant difference between the two mental states. The mental task of visualising pushing an object into space compared to visualising a relaxed state or feeling relaxed required more mental strength to perform and here it is shown to have direct neural changes of suppression of the alpha and beta oscillations. Global cortical engagement was observed, with the neural changes occurring approximately 1.5 seconds after the onset of the cue, which could be attributed to the latency in conjuring the visualisation of an action. These findings are supported by previous studies on VI, that have found alpha and beta suppression in the sensorimotor cortex (Lee et al., 2020) and the left temporal cortex (Castro et al., 2020), however presented with different stimuli cues.

Visualisation of action mental tasks elicited global cortical engagement which has implications of improving future VI based BCI applications, where visualisation of actions could improve the generalisation of neural correlates for classification in DL models. These findings are significant as it shows potential for VI to be a BCI paradigm of choice for novice users, due to the ability of neural discriminate changes being successfully performed.

To further address these findings and to address the BCI illiteracy problem within and between subject BCI performance variability, VVIQ scores were obtained. The hypothesis that those who scored higher on the VVIQ questionnaire would be able to perform the VI paradigm better than those who scored lower was supported. When the dataset was separated out by the high VVIQ scores and low VVIQ, only the high VVIQ group showed the significant alpha and beta suppression in the “push” condition compared to the “relax” condition. To the author’s knowledge this is the first study to assess the vividness capabilities of the novice users with the VI based BCI. However, the cluster-based permutation test between the two groups was not able to be performed due to the unequal sample sizes of the high VVIQ group to low VVIQ group. This finding suggests that future research in utilising VI based BCI could incorporate VVIQ into their screening process to ensure that VI paradigm is the optimal paradigm for novice users. To further assess the performance of VI, screening for equal sample sizes in low VVIQ and high VVIQ could be implemented.

### 4.2 Imagined Speech BCI

For the IS paradigm, there was no discernible significant neural oscillations extracted by the TF analysis. The hypothesis that there would be gamma oscillations in the language and speech areas whilst performing the IS paradigm was not supported. The cluster-based permutation analysis did not show any significant difference between the “relax” and “push” conditions of neural frequency changes, from repetition of either word in the mind. It could be due to the words “push” and “relax” not being phonetically differentiated, leading to poor discrimination. The number of syllables and length of the word could possibly affect this IS paradigm (Nguyen et al., 2018), as both words here are relatively short in length with only one syllable difference. Using two words was very simple compared to Lee et al., (2020) where 13 words were utilised, which found that “hello” vs “pain” was the most confusing word combination in both IS and VI and “pain” vs “water” was best-discriminated combination for IS. They suggested that semantic meaning of the words may affect the decoding discrimination between the words, however their study did not compare real words to pseudowords. Here, however, the meaning of the words did not aid in discrimination, as the meaning of “push” and “relax” are quite different, but it may be crucial for future works to assess word selection for IS paradigms.

### 4.3 1D-CNN for Classification in VI and IS BCIs

The results presented here of the 1D-CNN demonstrated high-performance in predicting the two classes “push” and “relax” in both VI and IS paradigms. With the VI classification performance significantly outperforming the IS paradigm by approximately 11.6% accuracy. This could be attributed to the higher proportion of the dataset showing significant alpha and beta suppression during “push” and “relax” conditions. However, although the cluster-based permutation test did not show any significant neural changes in the IS paradigm, the 1D-CNN was able to classify between the two conditions with 77.7% performance accuracy. This could be attributed to the ability of the 1D-CNN to extract features during the five convolutional layers and classify in the remaining layers as opposed to TF extraction.

In addition, the use of batch normalisation during training, reLU activation, and random cropped training for data augmentation, boosted the classification performance of the 1D-CNN model. These results are reproducible with previous works (Mattioli et al., (2021), Schirrmeister et al., (2018)) who used similar hyperparameters in their DL models for decoding EEG signals for BCIs. During the network training to optimise the hyperparameters, in random cropping training the overlap factor was used to increase the dataset by overlapping the current data by replicating it, found that increasing it from 0.5 to 0.9 showed an increase from 50-99% percentage accuracy of the model. This data augmentation method was intended to increase the training dataset to avoid overfitting for the 1D-CNN, amongst other methods such as batch normalisation, early stopping and checkpointing. Despite the performance metrics demonstrating high-performance, upon assessing the loss and accuracy plots of the best and worst performing subjects in both VI and IS paradigms, it was apparent that the model had overfitted. The unstable nature of the validation loss and accuracy curves indicated uncertainty for the 1D-CNN to predict the validation set, suggesting that there is under-representation of the validation set compared to the training set. This may be due to the random cropped training applied to the training, validation and test datasets as opposed to data augmentation only applied to the training dataset in Mattioli et al., (2021). Increasing the trial per condition, per paradigm, per subject would be the prominent suggestion to avoid overfitting and validate the findings in this study.

The previous studies that have utilised CNN to decode covert speech (IS) were Cooney et al., (2019 & 2020) which achieved 34% and 35.68% accuracy, respectively, for classification of vowels with the same dataset of 15 subjects. Tamm et al., (2020) achieved 24% accuracy for classification of five vowels and six words utilising the same dataset in the two prior studies of 15 subjects. Whereas for decoding VI with CNN, Llorella et al., (2020) achieved 60% accuracy for classification of visual objects in 8 participants. Castro et al., (2020) utilised a CNN-LSTM which achieved 89.46% for classification of shapes with the dataset from Nieles et al., (2018) of 5 subjects, which does not seem plausible given the very small dataset. This current study performed well based on the size of the dataset at 20 subjects, and 1D-CNN performance metrics of 89.3% and 77.87% performance accuracy for VI and IS, respectively. Despite this, the model was still able to meet the high-performing accuracy metrics, which suggests this adapted 1D-CNN has some validity in classifying VI and IS for BCIs.

### 4.4 Limitations and future research

The limitations for each paradigm were mentioned in the relevant sections above and due to the time constraint and complexity of this study project, further exploration of optimising the training of the adapted 1D-CNN could not be performed. Future research should focus on increasing the trials per condition per paradigm in each subject, to further develop the 1D-CNN as a classification model for BCI applications. Rethinking the regions of interest as inputs into the 1D-CNN could improve generalisation and further develop commercial BCI headsets with smaller channels focused on significant relevant cortical regions. More questionnaires to assess cognitive function, personality types, behaviour could be implemented for the IS and VI paradigms. For future research into IS, it would be pivotal to include a rest class to compare the IS mentals tasks against, and thorough thought into the type of words chosen for better discrimination.

### 4.5 Conclusion

To conclude, the findings in this study of alpha and beta suppression during the “push” or mentally strenuous condition compared to “relax”, contributes to the current body of work on VI as a BCI paradigm. It suggests that VI BCI is a dynamic and plausible option compared to standard BCI paradigms, and that VI BCI illiteracy could potentially be controlled using VVIQ. The potential of the 1D-CNN model in classification of VI and IS BCIs is also demonstrated. This study took a user-centric approach to address the major challenges contributing to the development of reliable, accessible, and commercially viable BCI systems through addressing the underlying neural mechanisms, cognitive and behavioural factors, and current deep learning techniques.

## Acknowledgements

The author would like to acknowledge the research project was a partial fulfilment to a MSc in Computational Cognitive Neuroscience at Goldsmiths, University of London and was industry collaborative with LiquidWeb. The author sincerely thanks her supervisors Giuseppe Lai^1,2^, Maria Herrojo-Ruiz^1^, and David Landi^2^ for their insightful advice, guidance and support throughout the research process. Lastly, to her father for teaching her the value of hard work, to her mother for guidance of kindness and respect, to her sisters and friends for their limitless patience and encouragement when she needed it the most.

1 Department of Computing, Goldsmiths, University of London
2 LiquidWeb s.r.l.

\***Supplementary materials** (code scripts) can be found here: https://github.com/dia-tle

